# The asymmetric chemical structures of two mating pheromones reflect their differential roles in conjugation of *Schizosaccharomyces pombe*

**DOI:** 10.1101/548586

**Authors:** Taisuke Seike, Hiromi Maekawa, Taro Nakamura, Chikashi Shimoda

**Affiliations:** Microbial Genetics Laboratory, Genetic Strains Research Center, National Institute of Genetics, 1111 Yata, Mishima, Shizuoka, 411-8540, Japan; Yeast Genetic Resources Laboratory, Graduate School of Engineering, Osaka University, 1-6 Yamadaoka, Suita, Osaka, 565-0871, Japan; Department of Biology, Graduate School of Science, Osaka City University, 3-3-138 Sugimoto, Sumiyoshi-ku, Osaka, 558-8585, Japan

**Keywords:** fission yeast, mating pheromone, G-protein coupled receptor, autocrine, sexual agglutination, cell fusion

## Abstract

In the fission yeast *Schizosaccharomyces pombe*, the mating reaction is controlled by two mating pheromones, M-factor and P-factor, secreted by M- and P-type cells, respectively. M-factor is a C-terminally farnesylated lipid peptide, whereas P-factor is a simple peptide. To examine whether this chemical asymmetry in the two pheromones is essential for conjugation, we constructed a mating system in which either pheromone can stimulate both M- and P-cells, and examined whether the resulting autocrine strains can mate. Autocrine M-cells responding to M-factor successfully mated with P-factor-less P-cells, indicating that P-factor is not essential for conjugation; by contrast, autocrine P-cells responding to P-factor were unable to mate with M-factor-less M-cells. The sterility of the autocrine P-cells was completely recovered by expressing the M-factor receptor. These observations indicate that the different chemical characteristics of the two types of pheromone, a lipid and a simple peptide, are not essential; however, a lipid peptide is absolutely required for successful mating. Our findings allow us to propose a model of the differential roles of M-factor and P-factor in conjugation of *S. pombe*.

**Summary statement:** Lipid pheromone peptides secreted locally from one cell may be concentrated at the fusion site with an opposite mating-type cell, which then polarizes to enable successful conjugation in *S. pombe*.

## Introduction

Sexual reproduction accelerates evolution by increasing the diversity of the gene pool. The fission yeast *Schizosaccharomyces pombe* has two mating types (sexes): *h^+^* (Plus) and *h^−^* (Minus) (Gutz et al., 1974; Egel, 1989; Egel, 2004;). Under nitrogen starvation, two haploid cells of opposite mating-type mate to produce a diploid zygote (Egel, 1971). The mating reaction is controlled by pheromonal communication, as illustrated in Fig. 1A. M-factor, secreted by M-cells, is a C-terminally farnesylated and *o*-methylated nonapeptide (Davey, 1991; Davey, 1992), whereas P-factor, secreted by P-cells, is a simple peptide of 23 amino acids (Imai and Yamamoto, 1994). These pheromone peptides are accepted by a specific G-protein-coupled receptor (GPCR): Mam2 for P-factor (Kitamura and Shimoda, 1991) and Map3 for M-factor (Tanaka et al., 1993). Activation of the G-protein associated with the receptors, Gpa1, transmits signals through a MAP kinase cascade comprising Byr2 (MAPKKK), Byr1 (MAPKK), and Spk1 (MAPK), resulting in the transcription of pheromone-induced genes essential for mating (Obara et al., 1991; Xu et al., 1994; Barr et al., 1996). The pheromone signaling pathway downstream of the activated GPCRs is shared in both cell types.

**Fig. 1.**
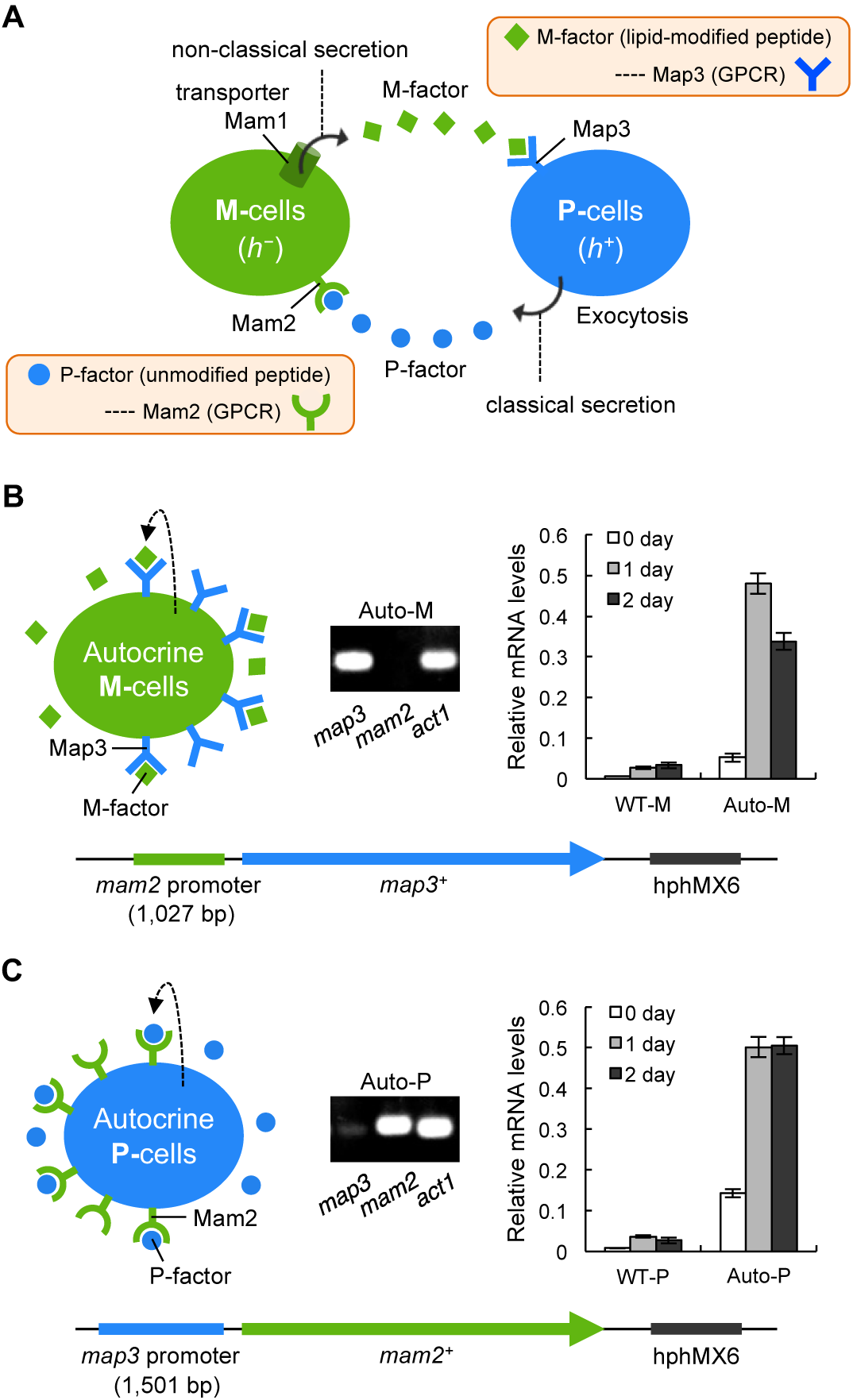
Strategy for creating autocrine haploid M- and P-strains. (A) Illustration of the mating pheromone signaling in *S. pombe*. (B) Construction of an autocrine haploid M-strain (FS600), in which Map3 is expressed by the *mam2* promoter (1,027 bp) in M-cells instead of native Mam2. (C) Construction of an autocrine haploid P-strain (FS683), in which Mam2 is expressed by the *map3* promoter (1,501 bp) in P-cells instead of native Map3. Quantitative RT-PCR shows ectopic mating-type-specific expression of *map3^+^* and *mam2^+^* in the respective autocrine M- and P-strains. The *act1^+^* gene was used as a control.

A key function of mating pheromones in yeasts is to guide a mating projection called a ‘shmoo’ (Moore et al., 2008). During mating, a cell senses a gradient of pheromones secreted by an opposite mating-type cell (Jackson and Hartwell, 1990; Segall, 1993), and forms a shmoo that is directed toward the center of the pheromone source. This polarized growth of *S. pombe* is regulated by the central regulator of cell polarity Cdc42, in complex with the guanine nucleotide exchange factor Scd1, and the scaffold protein Scd2 (Bendezú and Martin, 2013). Cells first adhere to opposite mating-type cells to form aggregates, which may help to stabilize the pheromone gradient, especially in liquid environments (Seike et al., 2013). Two mating-type-specific glycoproteins, Mam3 of M-cells (Xue-Franzén et al., 2006) and Map4 of P-cells (Sharifmoghadam et al., 2006; Xue-Franzén et al., 2006), are responsible for this sexual agglutination. Because cell fusion does not occur among cells lacking these proteins, even on solid medium (Sharifmoghadam et al., 2006; Seike et al., 2013), the close cell-to-cell contact mediated by Mam3 and Map4 is essential for cell fusion in *S. pombe*.

Pheromones in yeasts also play a role in the choice of a favorable mating partner. For example, cells of the budding yeast *Saccharomyces cerevisiae* choose a mating partner that produces the strongest pheromone signal (Jackson and Hartwell, 1990; Rogers and Greig, 2009). This is probably because the Cdc42 polarization complex forms at the highest concentration of pheromones from which the polarized growth starts (Bendezú and Martin, 2013). In fact, addition of exogenous pheromone to cells unable to produce their own pheromone does not rescue their ability to mate (Kjaerulff et al., 1994; Seike et al., 2013). Our previous study showed that the mating pheromones of *S. pombe* have a distal and proximal mode of action (Seike et al., 2013); that is, the general secretion of pheromones first induces sexual agglutination to increase cell density (‘distal’ action), and then locally secreted pheromones establish the polarity to influence mating partner choice (‘proximal’ action). Thus, mating steps regulated by these two different modes of pheromone action lead to successful conjugation.

In *S. pombe*, the pheromones for two mating types differ with respect to several properties: M-factor is a lipid-modified peptide (hydrophobic), and is specifically secreted by the ATP-binding cassette (ABC) transporter Mam1 (Christensen et al., 1997; Davey et al., 1997; Kjaerulff et al., 2005), whereas P-factor is an unmodified peptide (hydrophilic) that is secreted by exocytosis (Imai and Yamamoto, 1994) (see Fig. 1A). This chemical asymmetry between mating pheromone peptides is widely conserved across ascomycetes (Table S1) (Martin et al., 2011), although previous studies have suggested that pheromone asymmetry may not be required for mating in *S. cerevisiae* (Gonçalves-Sá and Murray, 2011). Furthermore, the biological significance of such asymmetric modifications of mating pheromones is not fully understood.

In this study, we have investigated the necessity of the chemical asymmetry of pheromone peptides in *S. pombe* by constructing autocrine cells that respond to their own secreted pheromone. We found that autocrine M-cells can mate with P-factor-less P-cells, whereas autocrine P-cells cannot mate with M-factor-less M-cells. Our findings clearly indicate that the chemical asymmetry of pheromones is not necessarily required for cell fusion, whereas a lipid peptide is essential for successful mating in *S. pombe*. We propose a model in which the hydrophobicity due to the farnesyl group on the pheromone plays a role in partner discrimination. Lipid peptide pheromones (i.e., M-factor) secreted locally from one cell might become concentrated near a cell of opposite mating type, resulting in successful conjugation.

## Results and Discussion

### Construction of haploid autocrine M- and P-strains

To investigate the necessity of the chemical asymmetry of the *S. pombe* mating pheromones, first, we attempted to construct a self-activated M-strain that responds to its own pheromone M-factor. The 5’-upstream sequence of the *mam2^+^* gene likely to contain the promoter was cloned into the plasmid pFA6a-hphMX6, which is used for chromosome integration (Fig. 1B). The activity of the promoter in M-type cells was confirmed by a *lacZ* fusion construct (Fig. S1). The pFA6a-hphMX6 (mam2^PRO^-map3) plasmid was integrated at the *map3* region of chromosome I of the heterothallic *mam2Δ* M-strain FS324. Transcription of the *map3^+^* gene in the resultant M-strain (FS600) was examined by a quantitative RT-PCR. As shown in Fig. 1B, the *map3* mRNA level markedly increased after 1 day of incubation of this M-strain in nitrogen-free medium (SSL−N), indicating that the M-factor receptor (*map3^+^*) was ectopically expressed. Therefore, we considered this self-activated M-strain to be an autocrine M-strain (Auto-M).

Next, we constructed a self-activated P-strain that responds to its own pheromone P-factor. To express the P-factor receptor gene (*mam2^+^*) in P-cells, the 5’-upstream sequence of the *map3^+^* gene likely to contain the promoter function was cloned into the plasmid pFA6a-hphMX6 (Fig. 1C). The map3^PRO^-*mam2*^+^ fusion gene was integrated in the authentic *mam2^+^* region on chromosome I of the heterothallic *map3Δ* P-strain FS618. In the resultant P-strain (FS683), an increase in *mam2* mRNA was confirmed by quantitative RT-PCR after 1 day of incubation in SSL−N medium (Fig. 1C). We thus concluded that an autocrine P-strain expressing only the P-factor receptor gene (*mam2^+^*) had been successfully constructed.

### Pheromone-induced polarized growth and sexual agglutinability in autocrine M- and P-cells

During mating, yeast cells sense a pheromone gradient and elongate a mating projection (shmoo) toward the source of the pheromone (Jackson and Hartwell, 1990). Extremely strong signals induce a default reaction, resulting in shmooing independent of a pheromone gradient (Bendezú and Martin, 2013). Therefore, we evaluated self-activation of the autocrine cells by their own secreted M-factor or P-factor by the observation of shmoo formation (marked elongation of polarized cells). On solid medium (MEA), the two autocrine haploid strains (FS600 and FS683) showed distinct shmoo formations (Fig. 2A), which were quantitatively assayed by measuring the ratio of cell length (L) to cell width (W) of individual cells (see Materials and methods). Whereas the L/W ratio in two wild-type haploid strains (FS324 and FS618) did not increase during 2 days of incubation, the L/W ratio of the autocrine haploid cells began to increase gradually after incubation started (Fig. 2A). Based on a definition of shmooing as an L/W ratio of more than 3.0 at 2 days, the autocrine M- and P-cells formed shmoos at a frequency of 26% and 44% (*n*=200 cells, each), respectively (Fig. 2B). In addition to shmooing, the autocrine M-cells were readily autolysed, as previously reported (Dudin et al., 2016), at a frequency of 39.3% ± 3.7% (*n*=1180), although lysis of the autocrine P-cells was not observed. This might be due to differences in the mechanisms of cellular pheromone signaling in M- and P-cells. Taken together, these data clearly demonstrated that the autocrine haploid cells were self-activated by their own pheromones.

**Fig. 2.**
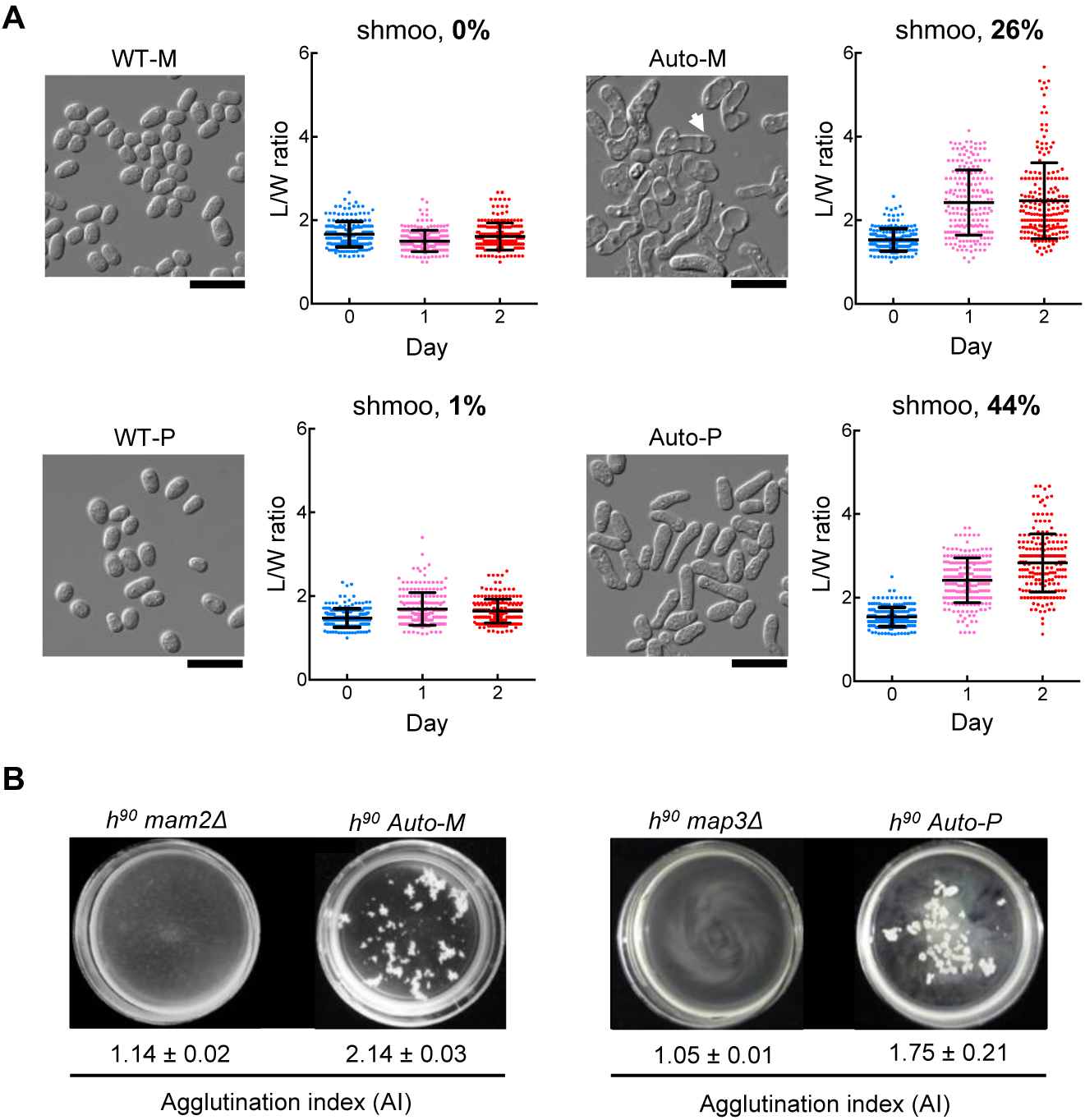
Polarized growth and sexual agglutination in autocrine M- and P-strains. (A) Cell morphology after induction of mating. Arrow shows an autolyzed cell. Scale bar, 10 µm. The ability for polarized growth was assessed by the length (L) to width (W) ratio of individual cells, and the frequency distribution of the L/W ratio is shown. Cells with a ratio of 3.0 or higher were defined as shmooing cells. For each strain, 200 cells were counted on each day of incubation. Data are given as the mean ± SD. (B) Measurement of the agglutination index (AI). Cultures in 3.5-cm petri dishes were gently shaken. The mean ± SD of triplicate samples is presented.

Conjugation of yeast cells commences after sexual agglutination in response to mating pheromones (Miyata et al., 1997; Seike et al., 2013) that is mediated by two mating-type-specific adhesin glycoproteins: Mam3 of M-cells and Map4 of P-cells (Sharifmoghadam et al., 2006; Xue-Franzén et al., 2006). We therefore examined sexual agglutinability in the autocrine M- and P-cells. We assayed agglutination intensity in two homothallic autocrine M- and P-strains (FS393 and TS167) cultured in SSL−N liquid medium. Two receptor-less homothallic strains (FS65 and TS160) were examined as a negative control. In both autocrine strains, strong agglutination was induced after 8 hours of incubation (Fig. 2B), suggesting that Mam3 and Map4 proteins were fully induced by their own pheromones at the cell surface. Our previous study showed that expression of the P-type-specific adhesin Map4 is completely dependent on pheromone signaling, whereas that of the M-type-specific Mam3 is induced only by starvation and further enhanced by pheromone treatment (Xue-Franzén et al., 2006; Seike et al., 2013). As expected, the receptor-less cells showed no agglutination (no visible aggregates, Fig. 2B). Collectively, these data indicated that both autocrine strains were fully self-activated, as judged by shmooing and sexual agglutination.

### Distinct differences in the mating ability of autocrine M- and P-cells

The mating competency of heterothallic autocrine M-cells (FS600) was examined by crossing with either wild-type heterothallic P-cells (FS127) or heterothallic P-factor-less P-cells (FS601). As shown in Fig. 3A and 3B, the autocrine M-cells completed conjugation with both wild-type and P-factor-less P-cells, although the mating frequency was significantly lower than that observed between wild-type pairs (40.73% ± 3.8% vs 14.7% ± 2.6%, Fig. 3B and Table S2), consistent with previous observations (Kitamura et al., 1996; Dudin et al., 2016). This result indicated that the asymmetry in the chemical nature of the two different mating pheromones is not essential for mating because *S. pombe* cells mated by using only M-factor.

**Fig. 3.**
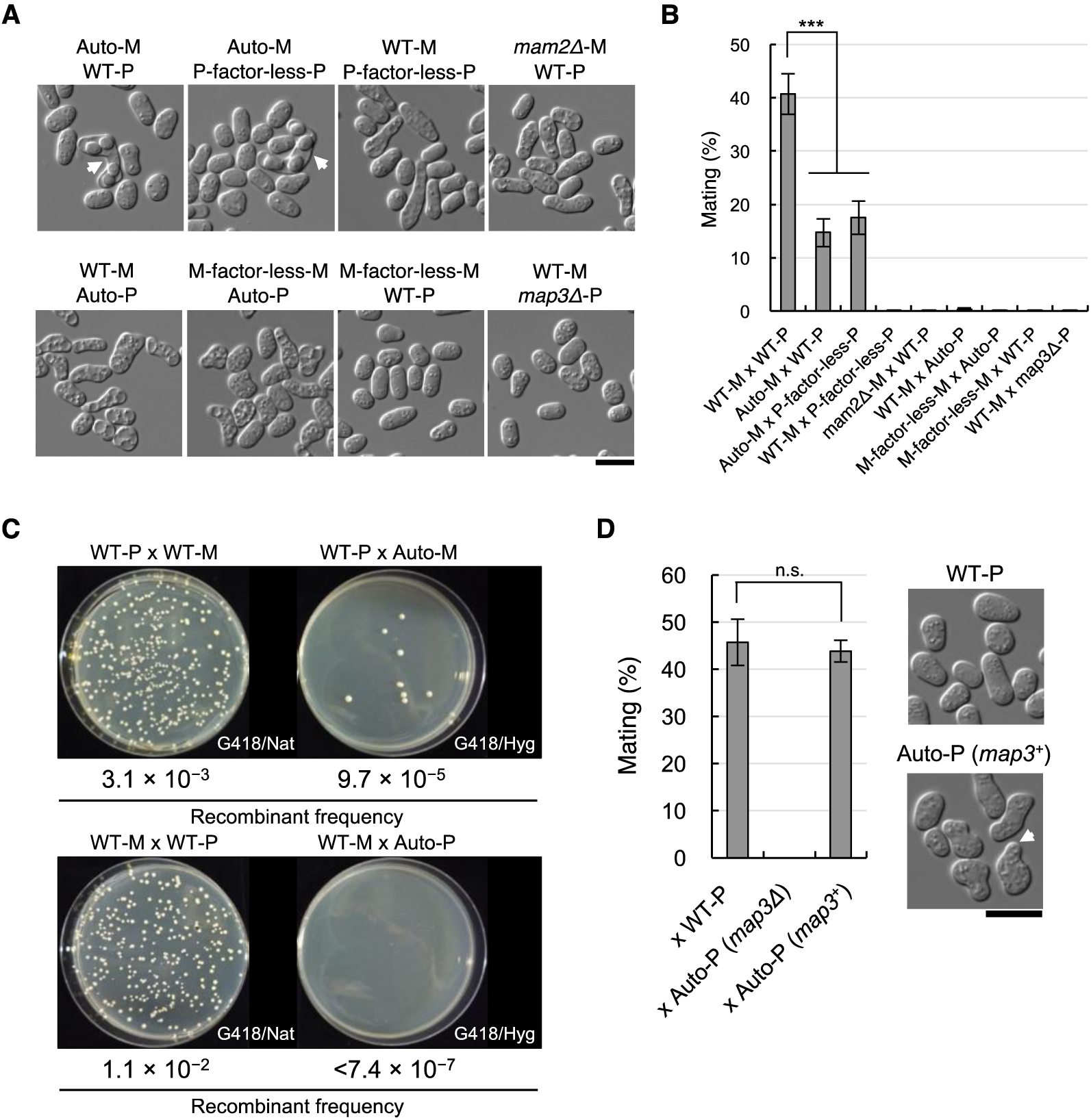
Mating ability of autocrine M- and P-strains. (A) Morphology of asci (arrows) in various combinations of two haploid strains. Scale bar, 5 µm. (B) Summary of the mating ability between two haploid strains seen in (A). Mating (%) was determined by measuring the frequency of zygotes. Data are the mean ± SD (*n*>300, each). \*\*\**P*<0.001. The exact percentages of mating are reported in Table S2. (C) Recombinant frequency of wild-type and autocrine strains. Heterothallic haploid strains were differentially marked by kanMX6, hphMX6, or natMX6 drug-resistant markers. The mean ± SD of triplicate samples is presented. (D) Effects on mating frequency of the co-expression of Map3 and Mam2 in autocrine P-cells (TS159). Expression of *map3^+^* in the autocrine P-strain recovered mating ability (43.9% ± 2.3%) to a level comparable to that of the wild-type pair (45.7% ± 4.9%). Data are given as the mean ± SD (n>300, each); n.s., not significant; scale bar, 5 µm.

To examine more quantitatively the lower mating ability of autocrine M-cells as compared with wild-type M-cells, wild-type P-cells (FS405), wild-type M-cells (FS424), and autocrine M-cells (FS600) were mixed in SSL−N at a cell number ratio of 2:1:1. Each of the three strains was differentially tagged by a drug-resistance marker (see Materials and methods). The mixed cell suspension in SSL−N was incubated at 28°C with gentle shaking for 24 hours, aliquots were diluted and spread on YEA medium containing different combinations of Geneticin (G418), Nourseothricin (Nat) and Hygromycin B (Hyg), and then the double-resistant clones generated by conjugation were counted. As shown in Fig. 3C, the recombinant frequency between autocrine M-cells and wild-type P-cells was significantly lower than that of the wild-type combination; that is, the frequency was about one-thirtieth of that of wild-type pairs. It is possible that the low selectability of autocrine M-cells as a mating partner might be due to the fixed (“default”) location of the polarity complex at cell poles.

The mating ability of autocrine P-cells (FS683) was also tested. Unexpectedly, the autocrine P-cells did not mate with either wild-type or M-factor-less M-cells (Fig. 3A and 3B). Furthermore, the ability of the autocrine P-cells to mate with M-cells was not recovered by exogenously added synthetic P-factor (5 µM) (Table S2). The sterility of the autocrine P-cells was further confirmed by the lack of recombinant colonies in a mixed culture of autocrine P-cells (FS683), wild-type P-cells (FS504), and wild-type M-cells (FS703) (Fig. 3C). Lastly, no diploid zygotes were observed between autocrine M- and P-cells by microscopy (Table S2). Taken together, these experiments suggest that the ability of P-cells to undergo cell fusion is likely to require stimulation by M-factor, but not P-factor.

### Establishment of cell polarity during mating

Our above findings suggested that cell fusion requires the correct localization of secreted pheromones. Previously, Martin’s group reported that the Cdc42 forms dynamic zones of activity at distinct locations over time prior to cell fusion, and these zones contain Mam1 (Bendezú and Martin, 2013). We therefore observed the localization of fluorescence-tagged Mam1 and Map3 proteins during mating in the wild-type cells. As expected, the fluorescent signals were mostly localized at the conjugation tip [Mam1 at the contact site, 86% (*n=*44); Map3 at the contact site, 88% (*n=*24); Fig. S2], consistent with previous data (Dudin et al., 2016). To observe the localization of Mam1 in more detail, Mam1-3×GFP was expressed together with mCherry-Psy1, a marker of the forespore membrane (Nakamura et al., 2001). As a result, Mam1 signal was concentrated at the fusion site on the membrane [co-localization of GFP and mCherry signal, 100% (*n=*29); Fig. S2]. In addition, we noticed that the Mam2 receptor for P-factor was frequently localized at the conjugation tip (Mam2 at the contact site, 82% (*n=*27); Fig. S2). These observations suggest that the local concentration of transporter and receptors at the fusion site leads to secure conjugation between mating partners.

We therefore considered that, for mating competency, P-cells might require local activation of the Map3 receptor. To examine this hypothesis, we tested the effect of expressing Map3 in autocrine P-cells (TS159), in addition to the P-factor receptor Mam2. Notably, this strain mated with wild-type M-cells, at a level comparable to that of wild-type cell pairs (Fig. 3D). This result clearly indicated that the expression of Map3 in autocrine P-cells completely recovered mating ability, suggesting that local signaling through Map3 is essential at the contact site between M- and P-cells. The cell polarity of P-cells is probably established by receiving M-factor secreted locally by M-cells. The haploid P-strain (TS159) co-expressing both Map3 and Mam2 produced a shorter shmoo as compared with autocrine haploid P-cells (FS683) (L/W ratio: 2.8 ± 0.7 vs 1.9 ± 0.4, Fig. 2B and 3D), probably because the two receptors compete for the same G protein, Gpa1, within the cell.

### Working hypothesis of conjugation in *S. pombe*

On the basis of the above experimental data, we propose the following model of the mating process in *S. pombe* (Fig. 4). In the first step, pheromone peptides are secreted in response to an environmental cue, such as nutritional starvation (Stage I). The hydrophilic simple peptide, P-factor, is easily diffused into the surrounding medium, and thus it will reach far-away cells, enabling those cells to become rapidly aware of the existence of favorable mating partners. Next, the relatively low concentrations of pheromones secreted into the milieu enhance the production of agglutinin for cell-to-cell contact, resulting in cell agglutination (Stage II). This physical contact between cells raises local pheromone concentrations at the cell surface, which fixes the active cell polarity complex comprising Cdc42 at the contact site (Bendezú and Martin, 2013). The M-factor transporter Mam1 is localized at the polarization site in M-cells, and simultaneously the M-factor receptor Map3 is located (or locally activated) near Mam1 in P-cells; thus, M-factor might be mainly secreted at the contact site. The hydrophobic lipid peptide, M-factor, is thought to be less diffused into the surrounding medium, and thus its concentration might be relatively high near P-cells. This local concentration of M-factor establishes the polarity of P-cells (Stage III), and polarized cell growth is induced. Cells of both mating types are firmly paired at the conjugation tip, and ultimately a pair of cells fuse to form a zygote. During the mating process, the localized secretion of M-factor is a critical step. The essential role of M-factor is dependent on the localization of Mam1 and on the retention of hydrophobic M-factor at the secretion site.

**Fig. 4.**
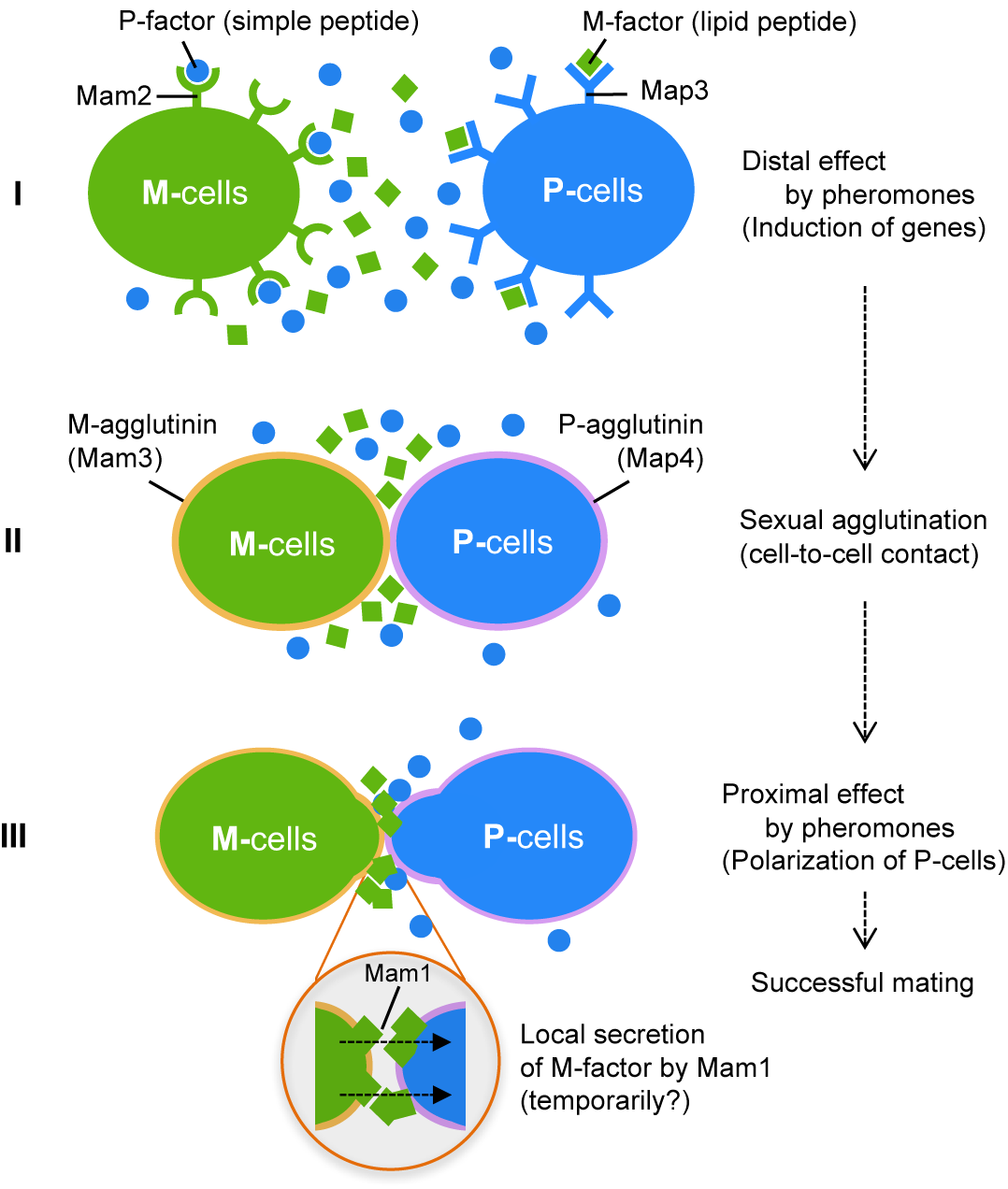
Working hypothesis of conjugation in *S. pombe*. The chemical asymmetry in the two mating pheromones reflects their differential roles in conjugation of *S. pombe*. In Stage I, uniform secretion of pheromone peptides facilitates the expression of pheromone-inducible genes, such as mating-type-specific agglutinins, Mam3, and Map4 (distal effect). In Stage II, pheromones induce sexual agglutinability in both M- and P-cells, which leads to stable cell-to-cell contact. In Stage III, polarized growth occurs at the contact site after agglutination. Hydrophobic M-factors are temporarily concentrated around P-cells, and these locally secreted M-factors establish the polarity of P-cells (proximal effect), leading to successful mating.

Our recent study in wild *S. pombe* strains demonstrated that M-factor and Map3 operate extremely stringently, whereas P-factor and Mam2 can tolerate variations in their structures to some extent (Seike et al., 2019), suggesting that M-factor communication plays an important role in defining the species. Therefore, the lipid peptide with a farnesyl group is likely to contribute to partner discrimination and cell fusion. We speculate that the hydrophobicity of pheromone peptides is probably beneficial for ascomycetes such as *S. pombe* living in a semi-liquid environment, enabling them to mate with an appropriate partner.

## Materials and methods

### Strains, media, and culture conditions

The *S. pombe* strains used in this study are listed in Table S3. Standard methods were used for growth, transformation, and genetic manipulation (Moreno et al., 1991). *S. pombe* cells were vegetatively grown in yeast extract (YE) medium supplemented with adenine sulfate (75 mg/l), uracil (50 mg/l), and leucine (50 mg/l). For solid medium, 15 g/l of Bacto Agar (BD Bioscience, Sparks, USA) was added to YE medium (YEA). Antibiotics [G418 (Nacalai Tesque, Kyoto, Japan), Hygromycin B (Wako Pure Chemical Industries, Ltd., Osaka, Japan) and Nourseothricin (Cosmo Bio Co., Ltd., Tokyo, Japan)] were added to YEA at a final concentration of 100 µg/ml. The solid medium used for mating was malt extract agar (MEA). To induce mating in liquid media, cells were shifted from SSL+N to nitrogen-free medium (SSL−N; minimal sporulation liquid medium without nitrogen) (Egel and Egel-Mitani, 1974; Gutz et al., 1974). Cells were incubated at 30°C for growth and at 28°C for mating, unless stated otherwise.

### RNA extraction followed by real-time PCR

Cells (OD_600_=0.1) were pre-cultured in YE liquid medium for 20 hours at 30°C. Cells at a density of 4×10^7^ cells/ml in 50-ml of SSL−N were continuously shaken at 28°C for 48 hours after sampling at the start point (0 hour). Next, 1 ml of culture was harvested for RNA extraction using an RNeasy^®^ Mini Kit (Qiagen, Hilden, Germany). DNA digestion was done to remove contaminating DNA in the solution before RNA clean up. For real-time PCR (RT-PCR), cDNA was synthesized from 500-ng samples of RNA using a SuperScript^®^ VILO cDNA Synthesis Kit (Invitrogen, Carlsbad, USA) according to experimental protocol. The program for RT-PCR was 94°C for 2 min, followed by 30 cycles of 98°C for 10 secs, 58°C for 30 secs, and 68°C for 30 secs. DNA segments containing *map3^+^*, *mam2^+^* and *act1^+^* (control) were amplified by using the following sets of primers: 5’-CTCCATGCCTGTTGTCTTTTGGTG-3’ / 5’-GCTGCCAAACATAGCAAGCGTAAG-3’; 5’-CTCCACTTAGCAGCAAGACCATTG-3’ / 5’-GCATCGGACCAAATGGAAATTGC-3’; and 5’-TGCTATCATGCGTCTTGATCTCGC-3’ / 5’-AAGCGACGTAGCAAAGTTTCTCC-3’, respectively.

### Observation of pheromone-induced cell growth (shmooing)

Cells were pre-cultured on a YEA plate at 30°C overnight, resuspended in sterilized water to a cell density of 1×10^8^ cells/ml, and 30-µl aliquots were spotted onto MEA plates, which were incubated for 1–2 days at 28°C. Cells were observed under a differential interference contrast (DIC) microscope, and photographs were taken. The length (L) and width (W) of individual cells were measured, and the L/W ratio was determined. In this study, cells with an L/W ratio of 3.0 or higher were defined as shmooing cells. In all experiments, at least 200 cells each were measured.

### Quantitative assay of sexual agglutination

The intensity of cell agglutination was photometrically determined as described previously (Seike et al., 2013). In brief, sexual agglutination was induced by shifting cells from SSL+N to SSL-N liquid medium at a density of 4×10^7^ cells/ml at 28°C. Cultures were incubated in 3.5-cm petri dishes (Thermo Fisher Scientific, Waltham, USA) with gentle shaking. The optical density at 600 nm (OD_600_) of an aliquot of the culture was measured, the cell suspension was then subjected to sonication (20 kHz) for 10 secs to disperse cell aggregates completely, and the OD_600_ was immediately measured again. The agglutination index (AI) was calculated by:

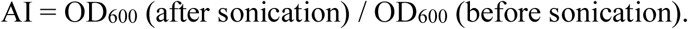

This value has been shown to be a function of the mean size of cell aggregates. In our experience, visible agglutination can be seen in samples with an AI of more than 1.1.

### Mating frequency between mating partners

Mating frequency was determined as described previously (Seike et al., 2012). Two haploid strains were separately grown on YEA plates overnight and then resuspended in sterilized water to a cell density of 1×10^8^ cells/ml. A 30-µl aliquot (15 µl of each strain) of cell suspension was then spotted onto a MEA plate, which was incubated for 1–2 days at 28°C. Cells were counted under a DIC microscope as described above. Cell types were classified into four groups: vegetative cells (V), zygotes (Z), asci (A), and free spores (S). The mating frequency was calculated according to the following equation:

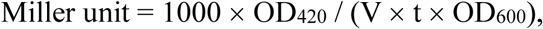

In general, triplicate samples (at least 1,000 cells) were counted, and the mean ± standard deviation (SD) was calculated.

### Quantitative assay of hybrid formation

The quantitative assay was done as described previously (Seike et al., 2013). Heterothallic haploid strains carrying a different drug-resistance marker (kanMX6, hphMX6, or natMX6) on their chromosomes (Seike et al., 2015) were mixed and then incubated in SSL−N medium for 2 days to induce mating. The cell suspension was diluted and spread on plates containing the appropriate combinations of drugs. The number of colonies was counted after 3 days of incubation. Three separate tests were carried out, and the mean ± SD was calculated as the recombinant frequency.

### Fluorescence imaging

Cells carrying Mam1-3×GFP, Map3-GFP, Mam2-GFP, or mCherry-Psy1 were immediately analysed by fluorescence microscopy without washing or fixation. Z-series images in 0.4-μm steps were captured with a DeltaVision system (Applied Precision, Issaquah, USA) equipped with GFP and TRITC filters (Chroma Technology Corp., Bellows Falls, USA), a 100×NA 1.4 UPlanSApo oil immersion objective (IX71; Olympus, Tokyo, Japan), and a camera (CoolSNAP HQ; Roper Scientific, Trenton, USA) and were quantified/processed with SoftWoRx 3.5.0 (Applied Precision, Issaquah, USA).

### Construction of a *lacZ* fusion gene and assay

A pDB248’-based multi-copy plasmid carrying the *mat1*-Pm/lacZ fusion gene (named pTA14) (Aono et al., 1994) was reconstructed to carry a *mam2* promoter-lacZ fusion construct. In brief, the *mat1*-Pm promoter sequence was replaced by an approximately 1-kb DNA fragment (Chr.I 4779661–4780687) containing the 5’-upstream sequence of the *mam2^+^* gene. The resulting plasmid pTA(mam2^PRO^-lacZ) was transformed into each of the heterothallic M- and P-strains (FS493 and FS494).

Enzyme activity was calculated as previously described (Miller, 1972). Cells were suspended in Z-buffer (60 mM Na_2_HPO_4_, 40 mM NaH_2_PO_4_, 10 mM KCl, 1 mM MgSO_4_・7H_2_O, and 50 mM β-mercaptoethanol) and stored at 80±C before the assay. Cells were permeabilized with chloroform and 0.1% sodium dodecyl sulfate. A solution of ONPG (*o*-nitrophenyl-β-galactoside) was added to the cell lysate and incubated at 28°C. The reaction mixture was centrifuged to remove cell debris and the optical density of the supernatant was determined at 420 nm (OD_420_). β-gal activity (Miller unit) was calculated according to the following equation:

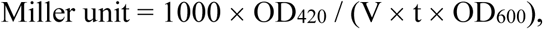

where v is the sample volume and t is the reaction time. The mean of triplicate samples was calculated.

### Statistical analysis

All experiments in this study were performed at least three times. The sample numbers used for statistical analysis are reported in the corresponding figure legends. A two-tailed unpaired Student’s *t*-test was used to evaluate the differences between strains. *P*-values are indicated in the figures (\*\*\**P*<0.001).

## Acknowledgements

We thank the National BioResource Project (NBRP), Japan, for providing yeast strains and plasmids.

## Author contributions

Conceptualization: T.S., C.S.; Methodology: T.S., H.M., C.S.; Validation: T.S., C.S.; Formal analysis: T.S., H.M., C.S.; Investigation: T.S., H.M., C.S.; Resources: T.S., H.M., T.N., C.S.; Data curation: T.S., C.S.; Writing – original draft: T.S.; Writing – review & editing: H.M., T.N., C.S.; Visualization: T.S., H.M., C.S.; Supervision: C.S.; Funding acquisition: T.S., T.N.

